# Preparation and Characterization of Inactivated Tick-Borne Encephalitis Virus Samples for Single Particle Imaging at European XFEL

**DOI:** 10.1101/2023.01.26.525647

**Authors:** Mikhail F. Vorovitch, Valeriya R Samygina, Evgeny Pichkur, Peter V Konarev, Georgy Peters, Evgeny V Khvatov, Alla L Ivanova, Ksenia K. Tuchynskaya, Olga I. Konyushko, Anton Y. Fedotov, Grigory Armeev, Konstantin V Shaytan, Filipe R N C Maia, Mikhail V. Kovalchuk, Dmitry I. Osolodkin, Aydar A. Ishmukhametov, Alexey M. Egorov

**Author notes:** Correspondence (V.R.S.); (A.M.E.). These authors contributed equally (Mikhail F. Vorovitch, Valeriya R Samygina).

## Abstract

X-ray imaging of virus particles at European XFEL could eventually allow solving their complete structure, potentially approaching resolution of other structural virology methods. To achieve this ambitious goal with today’s technologies, several mL of purified virus suspension containing at least 10^12^ particles per mL are required. Such large amounts of concentrated suspension have never before been obtained for enveloped viruses. Tick-borne encephalitis virus (TBEV) represents an attractive model system for the development of enveloped virus purification and concentration protocols, given the availability of large amounts of inactivated virus material provided by vaccine manufacturing facilities. Here we present the development of a TBEV vaccine purification and concentration scheme combined with a quality control protocol allowing substantial amounts of highly concentrated non-aggregated suspension to be obtained. Preliminary single particle imaging experiments were performed for this sample at European XFEL, showing distinct diffraction patterns.

## 1. Introduction

X-Ray Free Electron Laser (XFEL) is an extremely powerful instrument for a contemporary structural biologist (Fromme, 2015), and especially a structural virologist (Meents &Wiedorn, 2019). Hard X-ray FEL facilities (Emma *et al*., 2010; Ishikawa *et al*., 2012; Kang *et al*., 2017; Decking *et al*., 2020) are the brightest sources of X-ray radiation currently available, allowing the crystallographic imaging of biological objects at room temperature with femtosecond time resolution (Pandey *et al*., 2019) or single-particle imaging (SPI) (Miyashita & Joti, 2017; Meents &Wiedorn, 2019; Bielecki *et al*., 2020). These options are very attractive for structural virology, potentially allowing the structural mapping of different viral conformations, or virion dynamics (,,viral breathing” (Dowd & Pierson, 2018)), as well as detailed machinery behind the entry of viruses into cells (Rey & Lock, 2018). While the currently available gold standard methods of X-ray crystallography and cryo-electron microscopy mostly provide structural data for viruses at cryogenic conditions which may correspond to conformational spectrum may different from that present at physiological conditions(Ourmazd, 2019; Lyumkis, 2019). SPI at XFELs should eventually allow conformational sampling of viral particles and proteins near physiological temperature-in solution and eventually the virus-cell system. Such conformational analysis is already available for more strongly scattering, materials science samples (Ayyer *et al*., 2020; Zhuang *et al*., 2022). These data are of fundamental importance for the structure-based design of novel vaccines and antivirals.

Biological single particle analysis at XFELs is still in its early days and requires development of sample preparation methods, sample delivery systems, data processing and structure reconstruction algorithms, especially for such diverse objects as viruses (Meents &Wiedorn, 2019). Certain success was achieved with non-enveloped viruses, that form size-uniform well-structured particles or even crystals. For example, crystal structure was solved for bovine enterovirus 2 to the resolution of 2.3 Å using serial femtosecond crystallography (SFX) approaches (Roedig *et al*., 2017). Single-particle reconstruction of a mimivirus (capsid diameter 450 nm) electron density to the resolution of 125 nm was possible in early experiments at LCLS with injection via a gas dynamic virtual nozzle (GDVN) (Ekeberg *et al*., 2015), while the smaller Melbournevirus with capsid diameter 230 nm could be reconstructed with 28 nm resolution (Lundholm *et al*., 2018). Rice dwarf virus and bacteriophage PR772 (capsid diameters 70 nm in both cases, studied in the same setting) could be solved to a resolution of 13.5 and 7.8 nm, respectively (Kurta *et al*., 2017; Rose *et al*., 2018]. The PR772 dataset was used to explore the conformational changes that could correspond to the genome release from the viral particle (Hosseinizadeh *et al*., 2017). Proof-of-concept experiments were performed with mimivirus at the European XFEL, showing the possibility of single-particle imaging with MHz pulse frequency (Sobolev *et al*., 2020). Further development of single-particle virology at XFELs depends on the availability of high-quality samples of different viruses, well-characterised by other methods and possessing specific features of the structure.

Tick-borne encephalitis (TBE) is a severe viral disease that has a substantial epidemiological importance for Europe and Russia. The virus is transmitted through the bite of an infected tick. Clinical manifestations of TBE vary from uncomplicated fevers to potentially lethal encephalitis or meningoencephalitis. A chronic TBEV infection may manifest with lifelong neurological symptoms. Taken together, these factors negatively affect the quality of life in the endemic regions and require preventive and protective measures. Several widely used inactivated vaccines were developed to prevent TBE, and drug discovery programs are currently active (Ruzek *et al*., 2019). Despite a long (at least half-century) story of clinical use of these vaccines, they are still poorly characterised from a structural point of view. On the other hand, inactivated vaccines are produced in large volumes and may serve as a source of large amounts of viral particles required for XFEL studies. Structural studies of inactivated viruses are scarce in general and do not show substantial differences between the structure of intact and formaldehyde inactivated particles due to insufficient resolution and symmetry (Wang *at el*., 2012). A formaldehyde inactivated TBEV vaccine, usually referred to as “TBE vaccine Moscow” in the scientific literature, is manufactured in large volumes in FSASI “Chumakov FSC R&D IBP RAS” (Institute of Poliomyelitis) and widely used in clinical practice since the 1960’s (Vorovitch *et al*., 2015). Thus, TBE vaccines are considered as attractive model systems for SPI methodology development at XFELs.

Tick-borne encephalitis virus (TBEV) is a typical member of genus *Flavivirus* (family *Flaviviridae*), forming roughly spherical enveloped virions of about 50 nm diameter (Füzik *et al*., 2018) (Fig. 1). Five subtypes of the virus differ in antigen specificity and common disease manifestations; among the subtypes, European, Siberian, and Far-Eastern are the most studied (Ruzek *et al*., 2019). Particularly, TBE vaccine Moscow is derived from the Far-Eastern subtype strain Sofjin (Vorovitch *et al*., 2015). The structure of the outer protein shell of TBEV virions (strain Neudoerfl) was described with atomic resolution using cryo-electron microscopy (cryo-EM) (Füzik *et al*., 2018; Pulkkinen *et al*.,2022), and a lot of reference structural information is additionally available for other flaviviruses, such as dengue virus, Zika virus, West Nile virus, Japanese encephalitis virus, etc. (Hasan *et al*., 2018). Virions of flaviviruses are heterogeneous depending on their maturation state: all the samples may simultaneously contain mature, immature, half-mature, and damaged particles (Dowd & Pierson, 2018; Hasan *et al*., 2018; Plevka *et al*., 2011). This diversity limits the choice of methods for determining the structure of flavivirus virions. Despite that the flavivirus envelope protein shell may be characterised as well-described, structure of the nucleocapsid is elusive, and knowledge about it is limited (Therkelsen *et al*., 2018).

**Figure 1.**
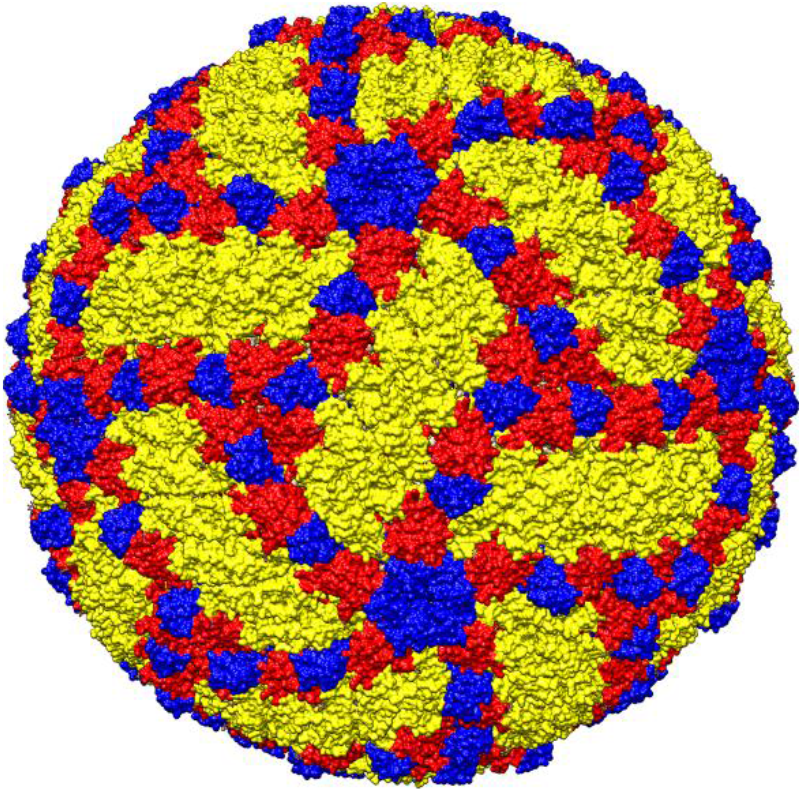
Overview of a mature TBEV virion structure obtained by cryo-electron microscopy (PDB ID 5O6A (Füzik *et al*., 2018). Envelope E protein is coloured by domains (domain I, red; domain II, yellow; domain III, blue).

The experimental method of SPI requires a substantial amount (several mL) of homogeneous, purified, and concentrated suspension of viral particles, containing at least 10^12^ (preferably 5×10^14^) particles per mL (Bielecki *et al*., 2019). Presently at European XFEL, it is required for pathogenic viruses to be inactivated. At the SPB/SFX instrument at European XFEL (Mancuso *et al*., 2019) the particles are delivered into the X-ray beam using an electrospray injector (Bielecki *et al*., 2019), so the sample solution needs to be sufficiently volatile. On the other hand, natural heterogeneity of enveloped virus particles may lead to the differential mobility of different particle types in the electric field (particularly, broken particles). Spontaneous aggregation of the viral particles is likely at high concentrations and may further obstruct the particle movement. To ensure the sample quality, the viral suspension needs to be characterised by different biochemical and physical methods, and biological relevance of the system needs to be confirmed. Maximization of content of mature particles, which are the most regular in form, in the sample is also desirable.

In this paper, we present the development of workflow for isolation, purification, concentration, and quality control approach for inactivated TBEV (iTBEV) particles from vaccine preparations. The quality control steps of this workflow employ a suite of virological, immunological, and physical methods. The obtained sample was successfully delivered to the X-ray interaction region point at the SPB/SFX instrument of European XFEL (Mancuso *et al*., 2019), and diffraction patterns were obtained from the individual iTBEV particles, thus providing a proof-of-concept for possibility of iTBEV single-particle imaging experiments.

## 2. Materials and Methods

### 2.1 Cells and Viruses

Prototype strain Sofjin-Chumakov of the Far-Eastern TBEV subtype (Genbank KC806252, Vorovitch *et al*., 2015), used in the manufacturing of inactivated TBEV vaccines at the FSASI “Chumakov FSC R&D IBP RAS” (Institute of Poliomyelitis), was used throughout the study. Virus was cultivated in the primary chicken embryo fibroblasts under the factory conditions. Briefly, cells, cultivated in 0.3% lactalbumine hydrolysate with Earl’s salt medium (FSASI “Chumakov FSC R&D IBP RAS” (Institute of Poliomyelitis), Russia) with 5% fetal bovine serum (FBS) (Biolot, Russia) and 50 μg/mL gentamicin at 37 °C, were infected with TBEV (MOI from 0.3 to 1 PFU/cell). Eagle MEM with 2% FBS, 2 mM L-glutamine, and 50 μg/mL gentamicin, was used as the support medium. Incubation was continued at 37 °C and virus-containing liquid (VCL) was harvested after 48-72 h.p.i. upon achievement of 75% cytopathic effect (CPE). VCL was titrated using plaque assay in porcine embryo kidney (PEK) cells as previously described (Dueva *et al*., 2020]. VCL was inactivated by incubation with 0.02 % formaldehyde at 32 °C for 72 h. Inactivation was controlled *in vitro* in plaque assay by the absence of viral plaques.

### 2.2 iTBEV Purification

iTBEV sample preparation started from the inactivated VCL (iVCL). iVCL was clarified by filtration using Profile II Filter Cartridge with absolute retention rate 0.5 μm (Pall, USA) followed by precipitation with 5% protamine sulphate (Sigma-Aldrich, USA) solution in borate buffered saline (BBS) (pH 8.5, Panreac, Spain) to achieve the final protamine sulphate concentration of 5 mg/mL. After incubation for 1 h at 4 °C, the mixture was centrifugated for 30 min at 12,000 rpm and 4 °C (Beckman J2-21, USA), and iVCL supernatant was collected. When iVCL volume was more than 400 mL, the tangential flow filtration was performed before protamine sulphate precipitation, using Pellicon 2 cassette with Biomax 300 kDa Membrane (Merck, USA).

<u>Size exclusion chromatography (SEC)</u> was performed using ÄKTAprocess 6 mm Std system controlled by the Unicorn 7.0 software (GE Healthcare, Sweden). Sepharose 6 Fast Flow beads (GE Healthcare) were packed in a column (packed volume 10 L, diameter 140 mm, bed height 560 mm, Kronlab) to purify the sample from other contaminants. Sanitization of the resin was carried out with 1 M NaOH followed by washing with water and equilibrating it with the TN buffer (0.1 M Tris-HCl, 0.13 M NaCl, pH 7.8), which was used in further chromatography steps. One liter of concentrated iVCL was applied onto the column. The applied flow rate was 150 mL/min. The column temperature was not controlled (ambient temperature). Protein UV detection was carried out at λ = 280 nm. Flow-through fractions were collected and analyzed by ELISA. Purification targets identified by ELISA were combined and precipitated in TNE/5 buffer (20 mM Tris-HCl, 26 mM NaCl, 1 mM EDTA, pH 7.8) by centrifugation at 35,000 rpm for 3 h at 4 °C (L90K, USA). Final precipitate was dissolved again in TNE/5 buffer.

<u>Ion exchange chromatography (IEC)</u> was performed using an ÄKTA Purifier protein purification system controlled by the software Unicorn 3.10 (GE Healthcare, Sweden). CIMmultus QA 1 mL column (BIA Separations, Slovenia) was applied operating in bind-and-elute mode. Two buffer solutions were used for IEC: TN/5 buffer (20 mM Tris-HCl, 26 mM NaCl) as the mobile phase A and TN buffer (100 mM Tris-HCl, 2.3 M NaCl) as the mobile phase B, both of them at pH 7.8. These solutions were used in different proportions during step-gradient elution. Column regeneration and equilibration were performed according to the manufacturer’s recommendations. In the binding step, 2 mL of the concentrated iVCL sample was diluted with mobile phase A to a final volume of 10 mL and then injected onto the column at flow rate of 1 mL/min. Unbound substances were washed out by applying 15-20 mL of mobile phase A until the conductivity signal dropped to the baseline. Chromatographic separation was performed by step-gradient elution with increasing of NaCl concentration (0.11–1.06 M). For all chromatographic experiments flow rate was 1 mL/min and fractions were collected in 0.5 mL aliquots. Protein concentration during IEC was monitored by absorbance at 280 nm. The flow-through fractions were analyzed by ELISA, the target fractions were combined and sequentially diluted with TNE/5 buffer, precipitated by ultracentrifugation (35,000 rpm for 3 h at 4 ° C, Beckman-Coulter L90K, USA), and then redissolved in TNE/5 buffer.

<u>Two-step ultracentrifugation.</u> iVCL supernatant was centrifugated for 3 h at 25000 rpm and 4 °C (Beckman-Coulter L90K, USA). Precipitate was resuspended in the TNE buffer (0.1 M Tris-HCl, 0.13 M NaCl, 1 mM EDTA, pH 7.8), transferred onto stepwise sucrose density gradient 15-60% w/w on BBS with 0.25% glycine (pH 9.0), and centrifuged for 3.5 h at 35000 rpm and 4 °C (L90K, USA). At the last step of purification, target fractions were identified by ELISA and SDS-PAGE electrophoresis, combined and reprecipitated in TNE/5 buffer by centrifugation at 35000 rpm for 3 h at 4 °C (L90K, USA). Final precipitate was dissolved again in TNE/5 buffer. Buffer conductivity was measured on the built-in conductivity monitor of the chromatographic system AKTA Pure 150M (GE Healthcare). The sample was kept at 4 °C no more than 7 d, or stored at −70 °C.

### 2.3 qRT-PCR

Concentration of RNA-containing particles in the sample was quantified using qRT-PCR. Reverse transcription was performed with M-MLV RT (Promega, USA) according to the manufacturer protocol with the primer Pow-TBE-3’: 5’-AGCGGGTGTTTTTCCGAGTC-3’. Quantitative PCR was performed on DNA Engine Analyzer (BioRad, USA) with RT-RT-qPCR kit (Syntol, Russia) as described previously (Dueva *et al*., 2020).

### 2.4 ELISA

ELISA was performed with commercially available VectoTBEV-antigen kit (D-1154, Vector-Best, Russia) according to the manufacturer guidelines. Optical density was measured at 450 nm (Multiskan Sky Microplate Spectrophotometer, Thermo Fisher Scientific, USA). The amount of E protein in the sample was assessed using the calibration curve for the standard antigen preparation (Timofeev *et al*.,1987; Ljapustin & Vorovich, 2010). Calculations were performed using the embedded PC software tool of Thermo Scientific SkanIt. Amount of the viral particles was estimated with the assumptions of 180 E protein molecules per viral particle and protein MW of 53 kDa (Füzik *et al*., 2018).

### 2.5 Spectrophotometric analysis of TBEV concentration

The RNA and total protein concentrations were determined in 2.5 μL samples on a Nanodrop spectrophotometer (Thermo Fisher Scientific, USA) at 260 and 280 nm, respectively, using the incorporated software. RNA concentration was in the range of 0.54 to 0.61 μg/mL. Total protein concentration was in the range of 0.98-1.1 mg/mL. The concentration of viral particles was calculated from the measured RNA and protein concentrations, assuming 180 E protein molecules per particle and protein MW of 54 kDa (Trauchessec *et al*., 2019).

### 2.6 Electrophoresis and Western Blotting

Electrophoresis in denaturing conditions was performed in 10% SDS-PAGE (Mini-Protein Tetra System, BIO-RAD, USA) using the MW marker kit 10–200 kDa (Thermo Fisher Scientific, USA). Bovine serum albumin (BSA, Combithek) was used as the protein standard (0.3–2 μg/lane). After electrophoresis the gel was treated with the water solution F (10% acetic acid, 40% EtOH), then the proteins were stained with 0.22% Coumassie brilliant blue G-250 (Sigma, USA) diluted in the solution F. Gel was scanned in the gel-doc system GBOX-CHEMIXX6-E (SynGene, USA). Densitometry of the lanes was performed in ImageJ 1.52v (NIH, USA). The quantity of E protein in the sample was determined using the calibration curve in the BSA amount vs. optical density coordinates.

For the Western blotting, protein samples after electrophoresis were transferred to the nitrocellulose membrane (Hybond ECL, Amersham, USA) in the buffer solution containing 48 mM Tris-HCl, 39 mM glycine, 0.037% SDS (w/v), 20% EtOH (w/v) using the Mini Trans-Blot cell module (BIO-RAD, USA). Empty binding sites of the membrane were blocked with 5% defatted milk (Semper, Sweden) in PBS-T buffer (80 mM Na_2_HPO_4_, 20 mM NaH_2_PO_4_, 100 mM NaCl, 0.1% (w/v) Tween-20). Membrane was incubated with TBEV detector antibodies (rabbit polyclonal antibodies to EK-328 strain or mouse monoclonal antibodies 14D5 (Tsekhanovskaya, N.A. *et al*., 1993) for 1 h at 37 °C, washed with PBS-T buffer, supplied with peroxidase conjugates against rabbit or mouse IgG (Goat Anti-Rabbit IgG H&L (HRP), ab. 6721; Goat Anti-Mouse IgG H&L (HRP), ab. 6789; Abcam, UK), incubated for 1 h at 37 °C, and washed. Immune complexes were detected using chemiluminescent reagent ECL (Amersham, USA) and LucentBlue X-ray Film (Advapsta, USA).

### 2.7 Dynamic Light Scattering

The particle size distribution was studied using Dynamic Light Scattering (DLS) method on SpectroLight 600 (Xtal Concepts GmbH, Germany) at 20 °C in plastic 96-well microplates for 1.5 μL aliquots under the paraffin oil. Each aliquot was placed in 2 wells and measured 5 times with 20 s intervals. Alternatively, measurements were performed at the same temperature using DynaPro NanoStar (Wyatt Technology) in the disposable plastic cuvette. 5 μL aliquots of the sample were measured 20 times with 10 s intervals. *Hydrodynamic radii were calculated by the* firmware *of* instruments.

### 2.8 Nanoparticle tracking analysis

Nanoparticle tracking analysis (NTA) was performed using Malvern Panalytical NS300 equipped with a sample chamber with a 640-nm laser and a Viton fluoroelastomer O-ring. The samples were injected in the sample chamber with sterile syringes (BD Discardit II, New Jersey, USA) until the liquid reached the tip of the nozzle. All measurements were performed at room temperature. Sample was diluted in 15000 times in deionized water.

### 2.9 Differential Mobility Analysis

Differential mobility analysis (DMA) measurements were carried out in order to measure the particle size distribution after the sample was transferred into an aerosol. A scanning mobility particle sizer (SMPS) (TSI 3938) equipped with the TSI 3081A DMA, together with a TSI 3788 CPC particle counter, was used for the measurements. Using a sheath flow of 15 L/min we measured the size distribution of particles with mobility diameters between 6-217 nm divided into 100 logarithmically spaced size bins over a 50 s scan time.

### 2.10 Electron Microscopy

#### 2.10.1 Negative stain

Negative staining transmission electron microscopy (TEM) was performed using an FEI Titan transmission electron microscope, equipped with a CCD Gatan UltraScan 1000XP camera. 5 μL aliquot of the sample was applied to the freshly glow-discharged grids with continuous carbon film (TedPella, C only, 300 mesh). After 5 min excess sample solution was blotted out, followed by a fast application of uranyl acetate (UA). The grids with 1% UA solution were incubated for 1 min, the excess solution was removed and grids were dried before imaging at 300 kV. Images were collected at defocus values from 1 to 3 μm.

#### 2.10.2 CryoEM

For fast assessment of the sample’s quality with cryo-EM lacey grids (TedPella, C only) were glow discharged at 0.26 mBar, 30 mA for 30 s using PELCO easiGlow (TedPella). 3 μL aliquot of the sample was applied onto grids and vitrified using Vitrobot Mark IV at 4.5 °C, 100% humidity, 1.5 blot time. Grids were imaged in the cryo-TEM at 300 kV, using direct electron detector. Images were collected with the pixel size of 1.8 Å, defocus values from 1 to 2.5 μm. Movie processing and particle picking was performed in Warp (Tegunov & Cramer, 2019), followed by preliminary analysis in cryoSPARC (Punjiani *et al*., 2017) including 2D classification.

For high-resolution structure determination a sample of inactivated tick-borne encephalitis virus (prepared according to the optimised protocol) was used to prepare grids for cryo-EM. 3 μl of the sample was applied to a hydrophilized grids (Quantifoil, R1.2/1.3+2nm carbon) with an additional 2 nm layer of amorphous carbon. Grids were glow discharged prior to usage at 0.26 mBar, 15 mA for 12 s using PELCO easiGlow. Vitrification was carried out in a Vitrobot Mark IV (Thermo-Fisher Scientific) under standard conditions (humidity - 100%, T = 4.5°C, blot time - 3s). The grids were transferred to a cryogenic transmission electron microscope Krios G2 (Thermo-Fisher Scientific) equipped with a direct electron detector Falcon 2 and a Cs-corrector (CEOS). Data were obtained at an accelerating voltage of 300 kV, a pixel size of 0.863Å, defocuses in the range from −1.0 to −1.8 μm, and an electron flux of 40 e^-^/Å/s). A total of 2660 movies were collected, each consisting of 32 frames, with the total exposure time of 1.6 s.

Data pre-processing was performed on the fly using Warp. After motion correction, CTF estimation, and particle picking, 63,000 particles were extracted and imported into CryoSPARC, where the particles were binned to a box of 540 pixels. Two rounds of 2D classification resulted in a subset containing 44,000 particles. These particles were used for 3D refinement with aberration and defocus refinement and subsequent two rounds of 3D classification. The final iteration of 3D refinement followed by local refinement and reconstruction was performed using a subset of 37672 particles, resulting in a 3.02Å structure.

### 2.11 Small-Angle X-ray Scattering Measurements and Data Processing

Small-angle X-ray scattering (SAXS) measurements were performed for the iTBEV samples at the concentration range of 0.5 to 1.0 mg/mL (by total protein, 280 nm) in TNE/5 buffer (section 2.3).

Synchrotron radiation SAXS data were originally collected on the BioMUR beamline of the Kurchatov Synchrotron Radiation Source (National Research Centre “Kurchatov Institute”) (Peters *et al*., 2019). Measurements of the fresh inactivated TBEV samples from different purification protocols were performed in capillaries of 2.0 mm diameter at room temperature. Monochromatic radiation with a wavelength of 0.1445 nm (photon energy 8.1 keV) was used. The sample–detector distance for the measurements of iTBEV particles was 2.50 m, corresponding to the angular range of scattering vector s = 0.03÷1.1 nm^−1^, where s = 4πsin(q)/λ, 2q is the scattering angle, and λ is the X-ray wavelength. The SAXS data were recorded using a two-dimensional pixel detector PILATUS3 1M (DECTRIS, Switzerland). Exposure time was 10 minutes. The two-dimensional scattering pattern was then averaged over the radial direction in FIT2D (Hammersley, 2016). PRIMUS software (Konarev *et al*., 2003) was applied to subtract the contribution of buffer solution scattering from the scattering curves of virions.

Subsequent SAXS experiments were performed at the P12 beamline of the European Molecular Biology Laboratory (EMBL) at the PETRA III storage ring, DESY Hamburg (Blanchet *et al*., 2015). The suspensions of inactivated TBEV (obtained using the optimal purification protocol 3, as determined from the analysis of the preliminary SAXS experiments at BioMUR) were loaded using a robotic sample changer (Round *et al*., 2015) into a flow-through capillary of 1.7 mm diameter. The freshly prepared samples, as well as the ones stored at −80 °C for 5 months, were tested. The data were recorded using a Pilatus 6M detector (DECTRIS, Switzerland) with 20 x 0.05 seconds exposure time, at the sample to detector distance 3.10 m and wavelength 0.124 nm covering the momentum transfer range from 0.02 < *s* < 7.0 nm^-1^. The temperature was kept at 20 °C. The data collection and reduction were performed using BECQUEREL (Hajizadeh *et al*., 2018) and SASFLOW pipeline (Franke *et al*., 2012), including the comparison of frames for radiation damage, averaging, and buffer subtraction. The averaged frames were normalized to the transmitted beam using a beamstop with an integrated photodiode (Blanchet *et al*., 2015). No measurable radiation damage was detected by comparison of successive time frames.

The radius of gyration *R*_g_ of solute and the forward scattering *I*(0) were evaluated using the Guinier approximation (Guinier, 2017) at small angles (*s* < 1.3/*R*_g_) (assuming that the intensity is represented as *I*(*s*) = *I*(0) *exp*(−1/3(*sR*_g_)) and also from the entire scattering pattern in GNOM (Svergun, 1992) using the indirect Fourier transform. In the latter case, the distance distribution functions *p*(*r*) and the maximum particle dimensions *D*_max_ were also computed. The volume size distributions of TBEV particles within the approximation of polydisperse spheres were evaluated using the program MIXTURE (Konarev *et al*., 2003).

### 2.12 Single particle diffraction pattern processing and analysis

We use lit pixel counter (Hantke *et al*., 2014) in order to identify patterns with scattering above background. For each pattern, pixels with the signal higher on four standard deviations of the thermal noise than the dark signal are counted. Then the median and interquartile range (IQR) of the lit pixel numbers are computed for images taken in one train. Assuming the low hit rate the median and IQR are the robust estimators of the distribution of the lit pixel numbers for the images with the background signal. The images with the number of lit pixels higher median on more than 3.5 IQR are indicated as hits. We use cluster analysis in order to filter out anything except diffraction from single virion particle. We compute radial profiles from every diffraction pattern using pyFAI (Kieffer *et al*., 2020) and then we cluster them using hierarchical agglomerative clustering with cosine metric in log space. Bad clusters are excluded from the future analysis on the expert decision.

In order to estimate the size of the particles we fit the spherical form factor to the radial profile (Sobolev *et al*., 2020).

## 3. Results

### 3.1 iTBEV Isolation, Purification, and Concentration

The initial content of infectious virions in VCL was (3.4 ± 1.4)×10^8^ PFU/mL. Concentration of the viral particles in this VCL could not be measured directly due to presence of decoy particles, although could be roughly estimated as 10-100-fold higher than the infectious virion content (Dueva *et al*., 2020). After inactivation the absence of the infectious virus in the samples was confirmed *in vitro*. Initially, three different protocols of inactivated TBEV purification (Fig. 2) were compared in attempts to obtain highly purified viral particles using 100-400 mL of VCL. At the first step of all protocols, iVCL was clarified by filtration (Fig. 2, step AB1). On the next step protamine sulphate precipitation and centrifugation were performed. When the volume of iVCL was more than 400 mL, the tangential flow filtration (Fig. 2, step AB2) was performed before precipitation with protamine sulphate (Fig. 2, step AB3). In Protocol 1 size exclusion chromatography (SEC) was the next step. This Protocol is used in Sofjin strain TBEV vaccine production technology which includes steps AB1, AB2, AB3, B4 (Fig.2). Protocol 2 was modified from Protocol 1 with additional ion exchange chromatography step B5 which takes an additional 90 minutes. The main feature of Protocol 3 is ultracentrifugation usage. This protocol is the longest one, its duration is 8-9 hours longer than Protocol 1. At the last step of each protocol (step AB6, Fig. 2), target fractions were combined and reprecipitated in TNE/5 buffer, followed by centrifugation. Viral particle concentration in the final samples ranged from 2×10^11^ to 5×10^12^ mL^-1^ as estimated by ELISA and electrophoresis. iTBEV particle concentration for final samples prepared using Protocol 1 and Protocol 2 were 1.7×10^12^ and 2×10^11^, accordingly. Thus, the initial VCL was concentrated 3÷9×10^3^-fold.

**Figure 2.**
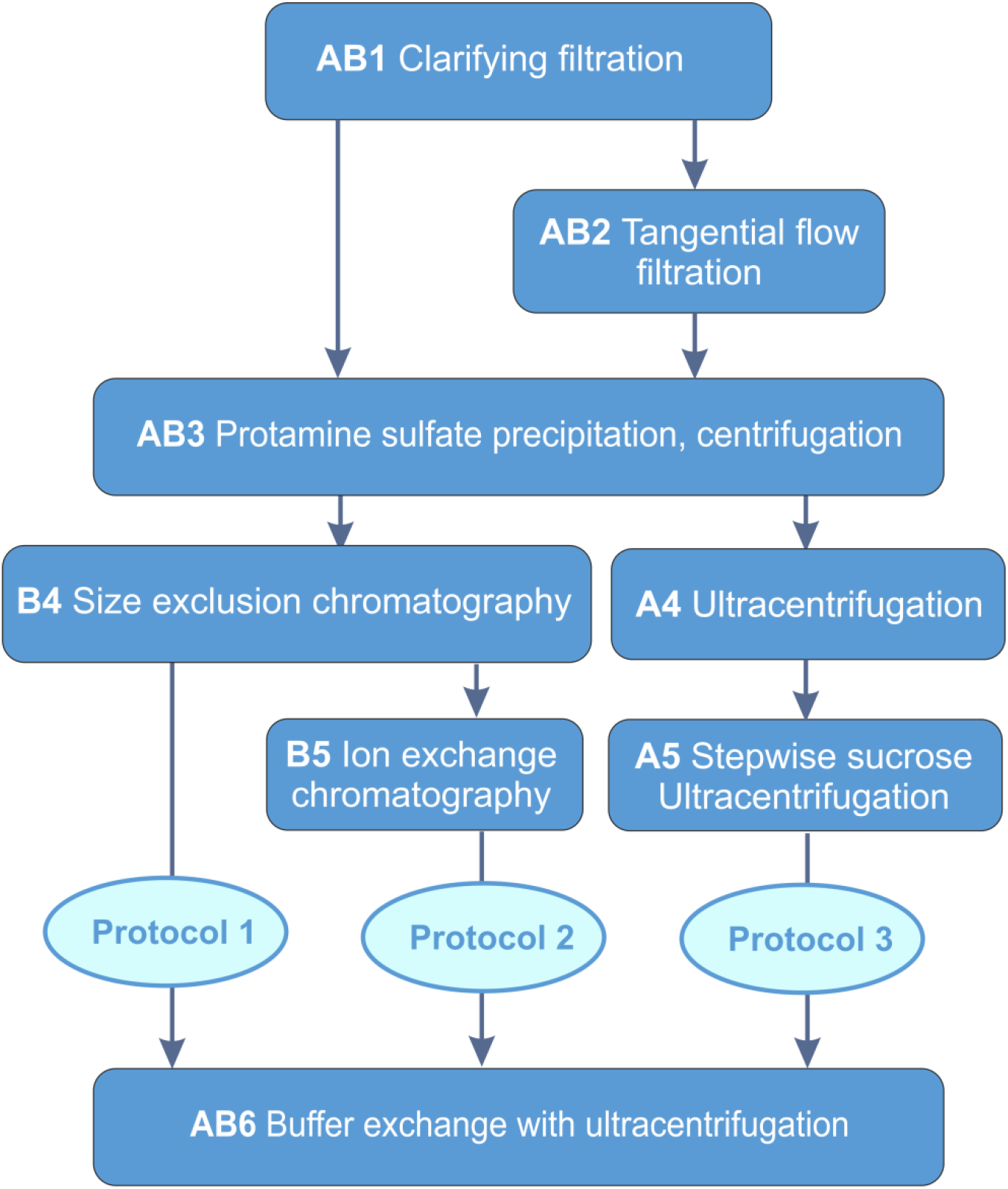
Scheme of TBEV purification protocols.

As soon as none of these techniques allowed direct measurement of the particle concentration and their intactness, TEM and SAXS were used to select the optimal protocol (see below). According to these data (see below), Protocol 3 gave the most consistent, homogeneous, pure, and concentrated suspension of viral particles.

### 3.2 Quality Control for iTBEV Samples and Choice of Optimal Purification Protocol

All three purification protocols gave high concentrations of iTBEV, but the biochemical characterization was not sufficient to confirm homogeneity and absence of particle aggregates. Thus, further characterization by physical methods was required to choose the optimal protocol, directly estimate the particle concentration, and ensure intactness of the particles.

#### 3.2.1 Electron microscopy

Transmission electron microscopy (TEM) was used for the visualization of the virions. Both negative stain electron microscopy and cryo-EM allowed estimating the size of virions, which was approximately 50 nm. The average diameter of 2D classes of the particles was estimated to be 47.1 nm (standard deviation 0.7 nm). Thus, the size of iTBEV particles was more uniform than in previous report (Morozova *et al*., 2014;). Both techniques allowed us to qualitatively confirm high particle concentration in the samples purified by different protocols. TEM was used as the main quality control technique in the initial development of purification protocols, allowing us to discard protocol 2 due to a loss of some sample at the step B5 (Fig 2) and an increase of the number of broken particles. Protocols 1 and 3 led to more concentrated and homogenous samples.

However, the negative stain requires the use of a contrasting agent (uranyl acetate), which may interact with the sample and produce some artificial assemblies (Fig. 3A). Cryo-EM images confirmed the size of iTBEV particles and the presence of several major types of particles, typical for flaviviruses, in the samples: mature, immature, semi-mature, and broken (Fig. S1, 3B, 4) (Füzik *et al*., 2018), however, some particles embedded in different layers of vitreous ice could not be distinguished from aggregates on the diagnostic images without the optimized ice thickness. Because both cryogenic and negative staining transmission microscopy rely on very small aliquots of the samples, the representativeness of the images may be insufficient, justifying the use of SAXS for further characterization.

**Figure 3.**
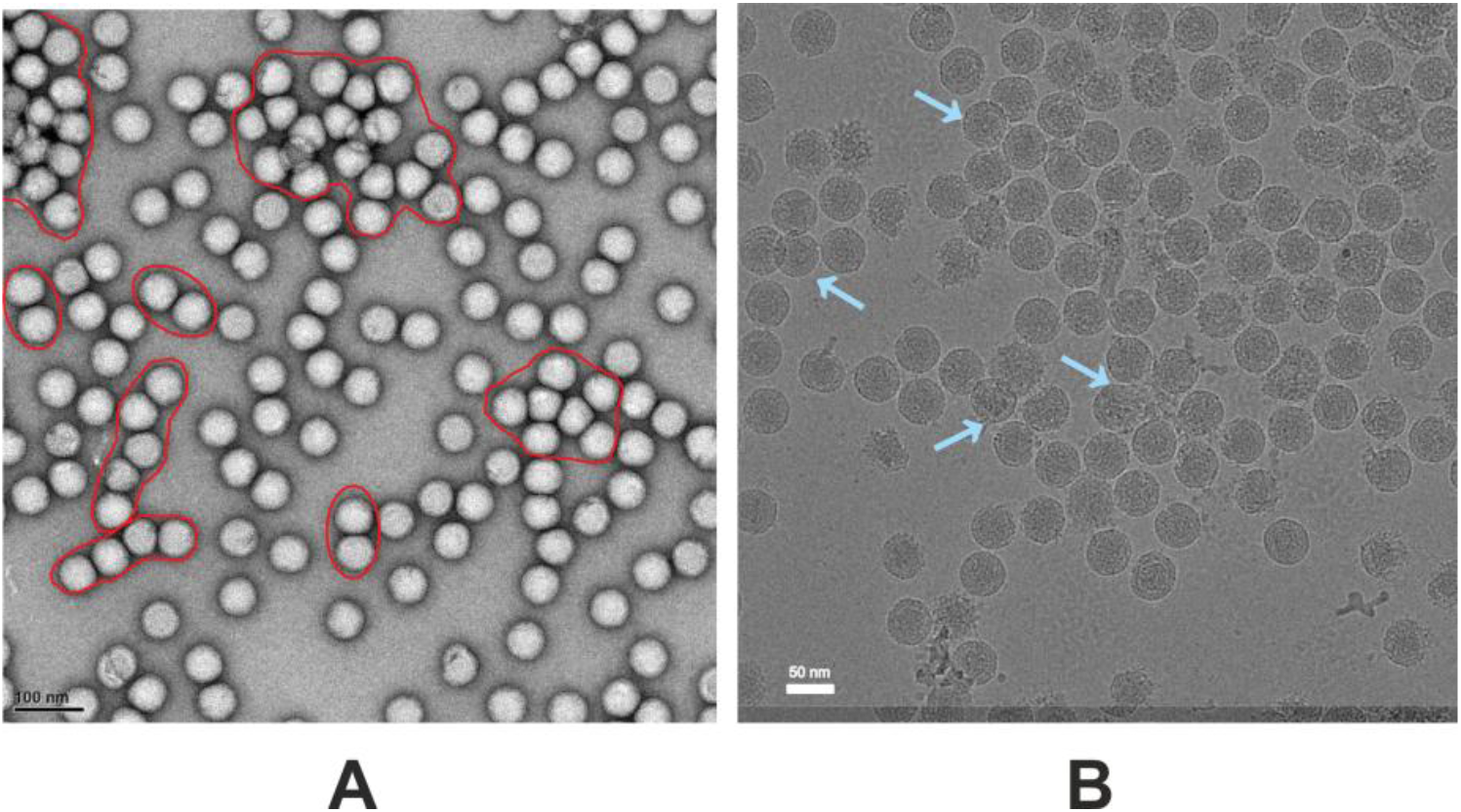
TEM of iTBEV. (**A)** Negative staining electron microscopy. Pseudo-aggregates are highlighted by red outline; (**B**) Cryo-electron microscopy of iTBEV. Pseudo-aggregates are indicated by blue arrows.

Moreover, we were able to solve the high-resolution iTBEV structure with the best resolution to date (3.02Å), further confirming the high quality of the sample obtained using protocol 3 (Fig.4, Fig.S4). A full size cryo-EM dataset, consisting of 2660 movies, demonstrates that the best samples contained only a low percentage of broken particles and very high percentage of mature particles based on 2D classification (Fig. 4C).

**Figure 4.**
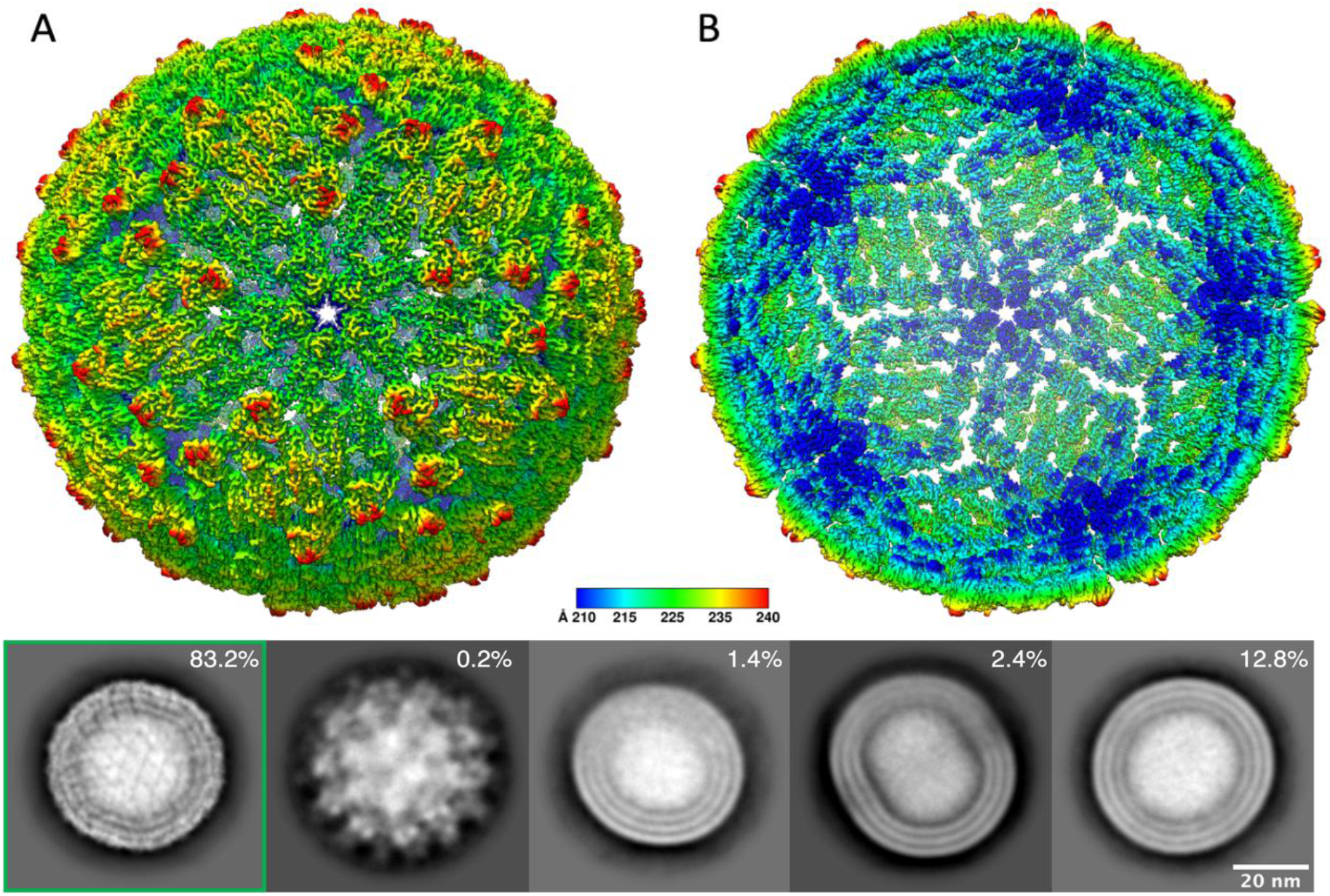
High-resolution cryoEM study using the developed protocol 3. (**A**) Radially colored virion shown in the direction of 5-fold symmetry. (**B**) Central cross-section of the cryo-EM structure. (**C**) Representative 2D classes of different particles in the high-resolution dataset and particle distribution across these classes. From left to right: mature, immature, half-mature, damaged, and elongated particles with broken symmetry. Particles marked with green were used for 3D structure determination.

#### 3.2.2 SAXS

We analyzed the experimental SAXS data from inactivated TBEV particles obtained using two different purification protocols, employing at the last step SEC (protocol 1) or ultracentrifugation (protocol 3). The radius of gyration and the maximal diameter of TBEV particles obtained using protocol 3 (Table 1) were consistent with the electron microscopy data. However, for the TBEV particles obtained with the protocol 1, these structural parameters were significantly higher (Table 1), indicating the presence of aggregated particles in the samples. Evaluation of volume size distribution within the approximation of polydisperse spheres showed that iTBEV particles obtained by protocol 3 have a narrow distribution with an average radius of 25 nm and a distribution width of 1 nm, while iTBEV particles obtained with protocols 1 are mixtures of spherical particles with the radii of 25 nm and 45 nm (Table 1). The experimental SAXS curves of iTBEV particles and the calculated distance distribution functions are displayed in Fig. 5A, the overall SAXS parameters are shown in Table 1.

**Figure 5.**
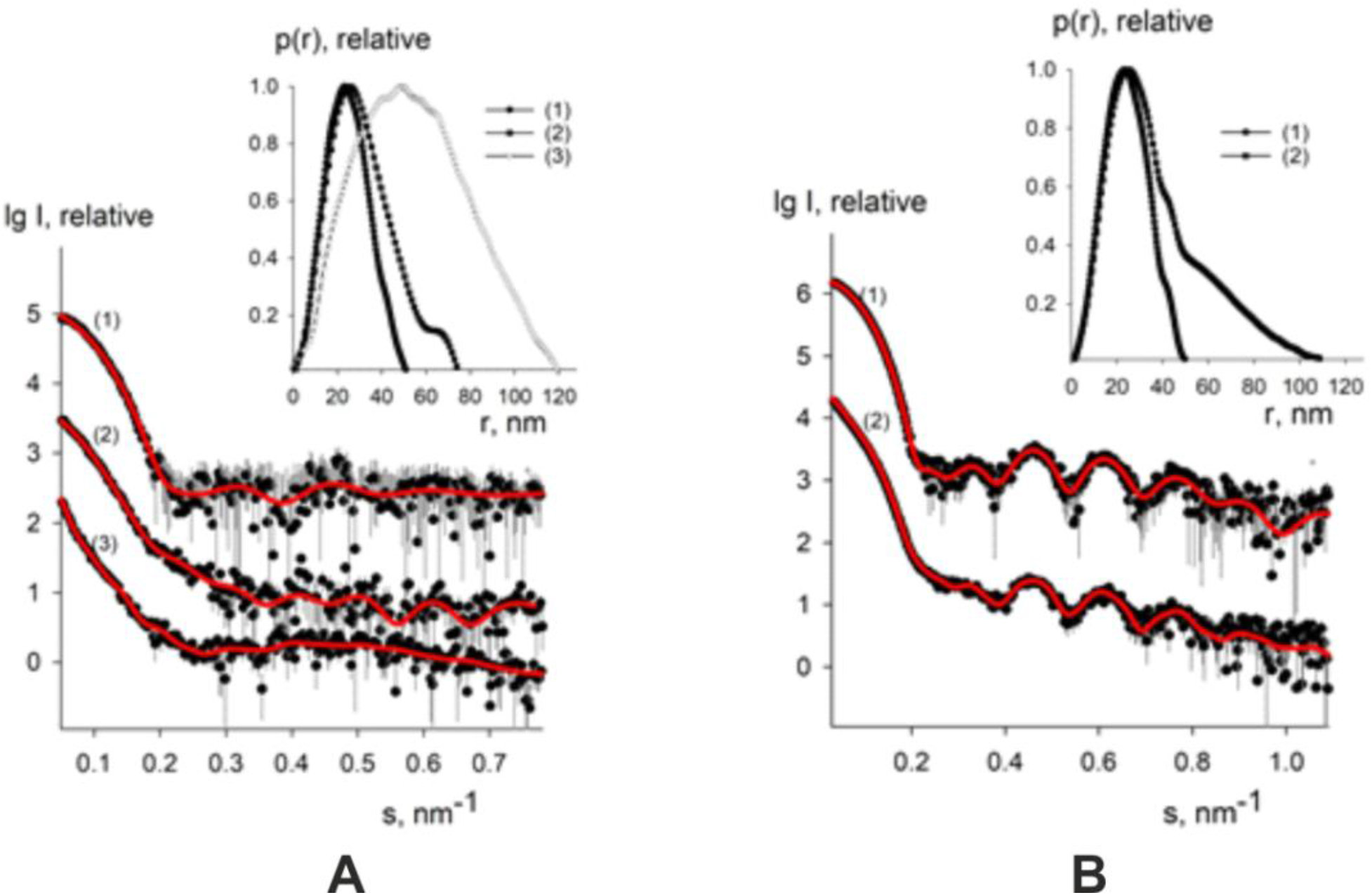
Small-angle X-ray scattering patterns from the iTBEV particles. (**A**) iTBEV data measured at BioMUR station obtained using three different purification protocols: curve (1) – purified using ultracentrifugation, curve (2) – purified using gel filtration (protocol 1), sample 2 (Table 1), curve (3) – purified using gel filtration (protocol 1), sample 3 (Table 1). (**B**) TBEV purified using ultracentrifugation, data measured at P12 station: curve (1) – from freshly prepared sample (protocol 3), curve (2) – from the sample stored at −80 °C during five months (sample 1). Experimental data are displayed as dots with error bars, scattering calculated by GNOM as solid red lines. The plots display the logarithm of the scattering intensity as a function of momentum transfer. The curves are displaced down by one logarithmic unit for clarity. The insets display the distance distribution functions estimated by GNOM. The curve numbering is the same as in the panel A.

**Table 1.** Overall parameters calculated from SAXS.

| Sample | Description | Beamline | $R_g^*$, nm | $D_{max}^{**}$, nm | Composition of volume size distribution |
| --- | --- | --- | --- | --- | --- |
| 1 | TBEV (protocol 3)<br>EuXFEL experiment<br>2145 | BioMUR,<br>KSRC | $18.7 \pm 0.2$ | $52.0 \pm 1.0$ | 100% of spheres with<br>$R = 25$ nm and the<br>width $dR = 1$ nm |
| 2 | TBEV (protocol 1) | BioMUR,<br>KSRC | $24.5 \pm 0.3$ | $75.0 \pm 2.0$ | 93% of spheres with<br>$R_1 = 25$ nm and the<br>width $dR_1 = 3$ nm<br>7% of spheres with<br>$R_2 = 45$ nm and the width $dR_2 = 6$ nm |
| 3 | TBEV (protocol 1) | BioMUR,<br>KSRC | $42.1 \pm 0.5$ | $120.0 \pm 3.0$ | 80% of spheres with<br>$R_1 = 25$ nm and the<br>width $dR_1 = 3$ nm<br>20% of spheres with<br>$R_2 = 45$ nm and the<br>width $dR_2 = 7$ nm |
| 4 | TBEV (protocol 3),<br>fresh<br>EuXFEL experiment<br>2316 | P12, Petra III | $18.4 \pm 0.2$ | $52.0 \pm 1.0$ | 100% of spheres with<br>$R = 25$ nm and the<br>width $dR = 1$ nm |
| 5 | TBEV (protocol 3)<br>(sample 1 stored<br>at -80 °C) | P12, Petra III | $30.3 \pm 0.4$ | $110.0 \pm 2.0$ | 90% of spheres with<br>$R_1 = 25$ nm and the<br>width $dR_1 = 3$ nm<br>10% of spheres with<br>$R_2 = 45$ nm and the<br>width $dR_2 = 5$ nm |

\* $R_g$ , radius of gyration; \*\* $D_{max}$ , maximum size of the particle. The composition of volume size distribution as estimated by MIXTURE.

Thus, it was found that the most suitable candidates for SPI experiments were the samples purified using ultracentrifugation in sucrose gradient (protocol 3).

#### 3.2.3 Large scale purification and biochemical characterization for SPI experiment

The sample prepared by protocol 3 was used for a nanofocussed SPI experiment 2316 on the SPB/SFX instrument of the European XFEL. The purification process was initiated from 10 L of inactivated VCL, which was purified by sequential cleaning and tangential flow filtration. On the next step protamine sulfate precipitation and centrifugation were performed, followed by the ultracentrifugation in the sucrose gradient. After these steps, virions were extracted from the sucrose and resuspended in TNE/5 buffer to give approx. 1 mL of suspension suitable for the sample storage and transportation (sample 745-11). This buffer was also considered acceptable for the electrospray injection of the sample (Bielecki *et al*., 2019), being diluted enough to avoid buffer aggregation and showing conductivity of 4.11 mSm/cm. Sample concentration estimates by different methods were in a good agreement (Table 2).

**Table 2.** iTBEV quantification in the sample 745-11 by different techniques.

| Technique | iTBEV particle concentration, mL<sup>-1</sup> |
| --- | --- |
| RNA photometry @ 260 nm | $\sim 3.1 \times 10^{12}$ |
| Protein E @ 280 nm | $\sim 9.8 \times 10^{12}$ |
| RT-PCR | $6.6 \times 10^{12}$ |
| ELISA | $2.3 \times 10^{12}$ |

SDS-PAGE was employed for purity assessment and identification of admixture proteins in the sample. 53 kDa protein was mostly present in the target fraction (Fig. 6A). Western-blotting with poly- and monoclonal E protein antibodies confirmed that this protein was the TBEV E protein (Fig. 6B). Pronounced antigenicity of the protein in Western blotting and ELISA confirms that at least some epitopes are preserved in the purified TBEV sample.

**Figure 6.**
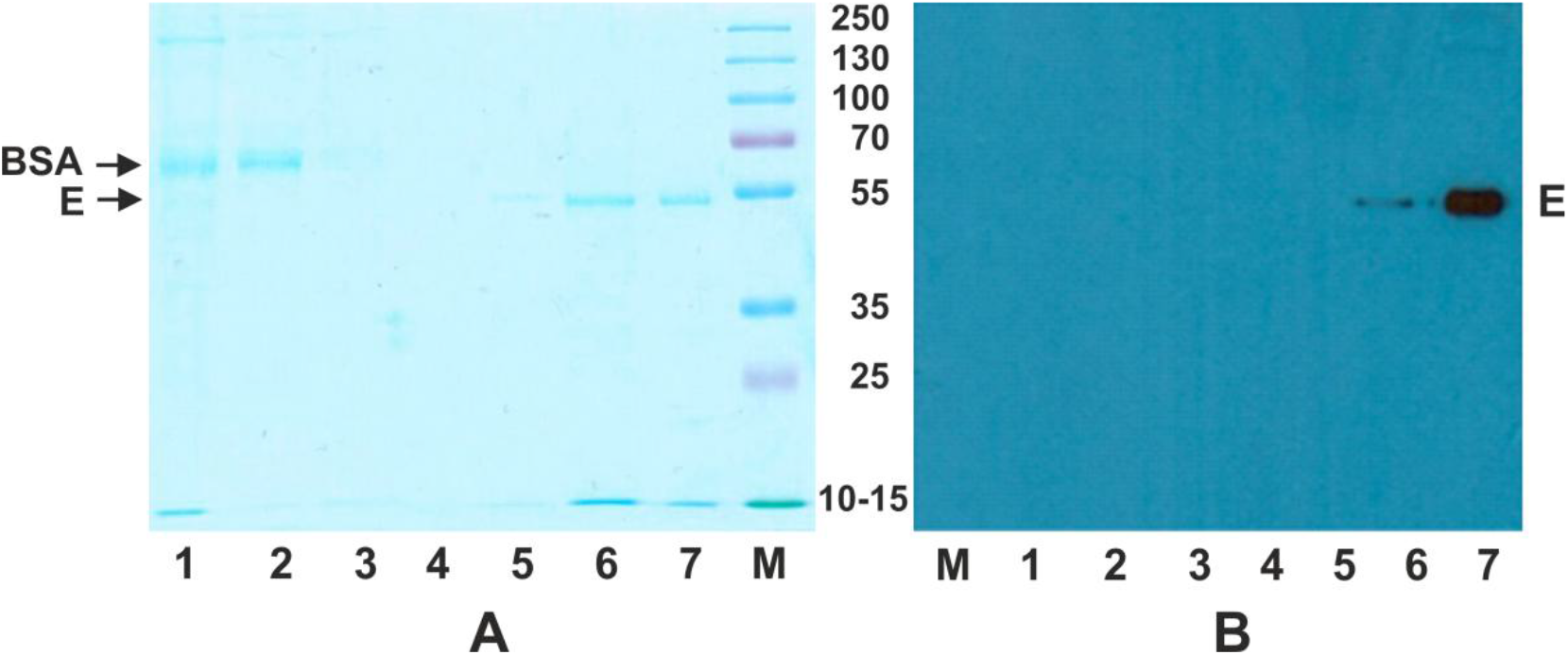
iTBEV sample 745-11 detection by electrophoresis and Western blotting. (**A**) 10% SDS-PAGE of initial preparation and sucrose gradient fractions after ultracentrifugation. Lane 1 – inactivated TBEV after centrifugation at 25000 rpm for 3h (BSA band at 67 kDa, E protein undetectable). Lanes 2-7 – sucrose gradient fractions after centrifugation at 35000 rpm for 3.5h; lanes 6 and 7 are target fractions; BSA is detectable only on lane 2. (**B**) Western blotting for sucrose gradient fractions. Lanes 1-6 – sucrose gradient fractions after centrifugation at 35000 rpm for 3.5h; lane 7 – merged target fractions after resuspending in the TNE/5 buffer. M - molecular weight markers.

### 3.3 On-site iTBEV characterization before experiment

The final assessment of the sample was performed prior to the beamtime to ensure there was no detrimental effect to the sample during transportation and also maximize sample quality for data collection at the SPB/SFX instrument. The iTBEV particle size and morphology were characterized using a variety of nanoparticle characterization techniques available at the XBI laboratory of the European XFEL (Han *et al*., 2021). Negative-stain transmission electron microscopy (TEM) (JEOL JEM2100-Plus) revealed a homogeneous population of particles with intact spherical morphology, indicating particles were not damaged during transportation at 4 °C (Fig. 7A). Complementary results were observed for iTBEV particles in solution with another direct imaging technique - atomic force microscopy (AFM) (JPK Nanowizard 4) (Fig. 7B). The sample was deposited on mica and AFM images were recorded in iTBEV buffer using force modulation mode. To characterize particles in solution dynamic light scattering (DLS) was used to determine the size distribution of iTBEV in the specimen. Being noninvasive, rapid and simple, it is useful for rapid assessment of the sample quality using small aliquots. A single peak was observed from two devices (WYATT Nanostar and Xtal concepts Spectrolight 600), corresponding to particles with a mean hydrodynamic radius of 26.4 ± 2.5 nm (Fig. 6C). The measured polydispersity did not exceed 18%, therefore the sample is considered monodisperse (Tomaszewska *et al*., 2013). Moreover, size information as well as particle concentration was determined by nanoparticle tracking analysis (NTA) (Malvern Panalytical NS300). The size distribution showed a major population with diameter 52.1 ± 3.1 nm and some minor peaks corresponding to aggregates (Fig. 7D). NTA measurements often require sample dilution, which may destroy or create new aggregates and affect the size distribution (Flipe *et al*., (2010)).These results corroborate the size distribution estimation by electron microscopy and SAXS, confirming the integrity and monodispersity of the virion suspension.

**Figure 7.**
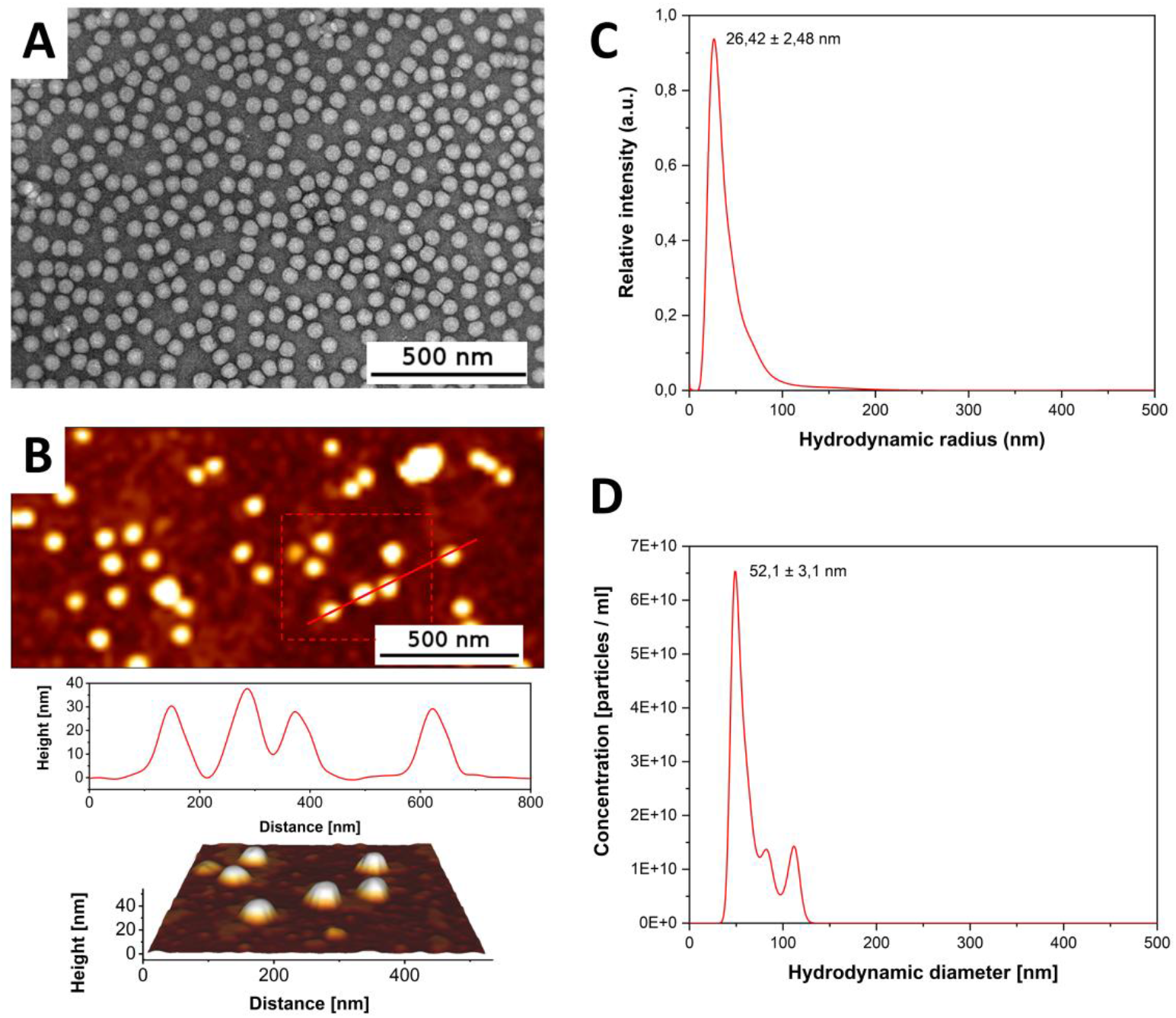
Characterization of iTBEV sample before beamtime. (**A**) TEM shows monodispersed spherical particles (scale bar is 500 nm). (**B**) AFM scan, line profile and 3D presentation of iTBEV nanoparticles. (**C**) DLS measurements of iTBEV particles reveal the hydrodynamic radius of the main sample fraction. (**D**) Hydrodynamic diameter as well as particle concentration is determined by NTA.

### 3.4 Electrospray Injection of TBEV particles

iTBEV particles were transferred into the gas phase via electrospray nebulization (Bielecki *et al*., 2019), resulting in droplets with diameter d = 145 nm with a liquid flowrate of 75 nL/min. For these conditions, the optimal sample concentration is approximately 5×10^14^ particles per mL based on maximizing the number of droplets with a single particle inside. The aerosolized particles were subsequently transported and focused into the X-ray interaction region with an aerodynamic lens stack (Liu *et al*., 1995; Hantke *et al*., 2018).

Directly prior to the X-ray experiment, monodispersity and stability of the aerosolized sample particles were investigated with a scanning mobility particle sizer (SMPS) (TSI 3938) equipped with the TSI 3080 DMA. Compared to solution measurements of the particle size distribution, the SMPS can be directly used to find conditions where the sample is efficiently transferred into the gas phase while minimizing the impact of non-volatile components in the sample buffer. In Figure 8 we show a SMPS trace of the iTBEV particles in the buffer used for the X-ray experiment. In contrast with the dynamic light scattering analysis, the size distribution consists of two main peaks at 11 and 45 nm respectively. The peak at 45 nm corresponds to the intact sample particle, while the 11 nm peak consists of salt that remains after the solution liquid have evaporated from empty droplets. Based on the size of the salt particle as well as the particle concentration ratio between salt and sample peak we can draw the following conclusions. First, the optimal sample concentration is approximately 4 times higher than what we have available in this sample batch. Secondly, the concentration of non-volatile salt in the solution buffer will give rise to a salt layer on the sample particle with a thickness of approximately 5 Å. In addition, two minor peaks at 30 and 59 nm respectively can be seen in the inset, where the same data is shown on a logarithmic scale. The 59 nm peak probably arises from instances where two iTBEV particles occupy the same droplet. The presence of this peak is a good sign that the sample concentration is close to optimal. The 30 nm peak on the other hand is likely due to iTBEV fragments or subviral particles.

**Figure 8.**
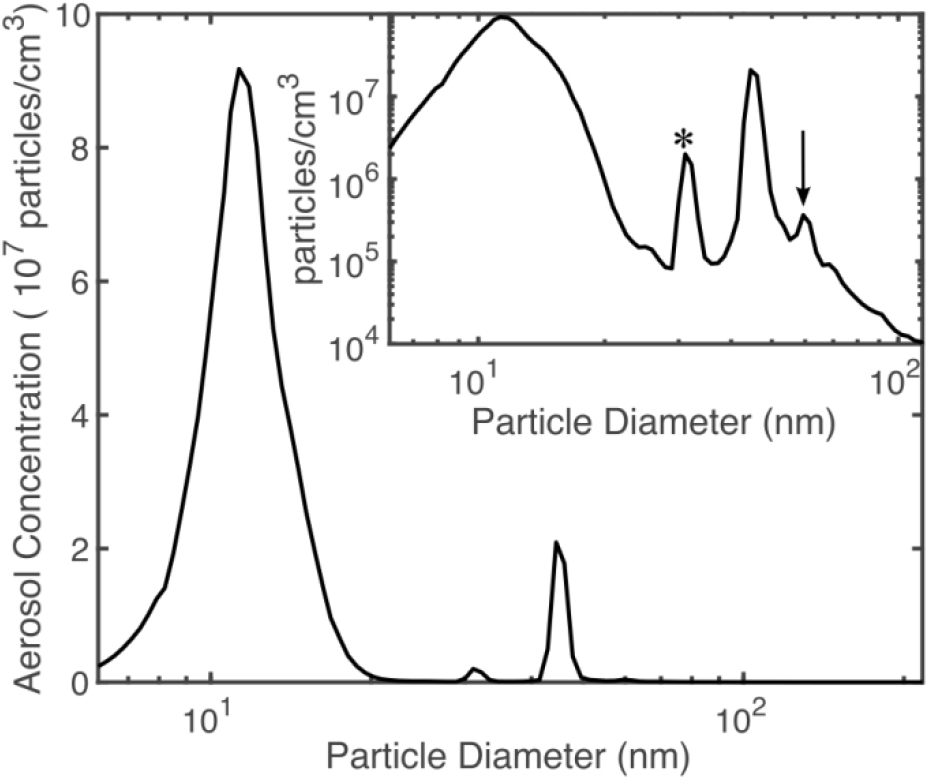
SMPS analysis of the iTBEV sample. The inset shows the data on a logarithmic scale, with the 30 nm fragments indicated with a star and the 59 nm iTBEV dimer indicated with an arrow.

### 3.5 Single Particle Diffraction of TBEV Virions

Diffraction patterns of single TBEV particles were collected at the SPB/SFX instrument at the European XFEL (Mancuso *et al*., 2019) as part of proposals 2145 and 2316. 6 keV X-ray pulses, each with 2.5 mJ pulse energy, were focused with a KB mirror pair to a spot with less than 500 nm FWHM. X-ray pulses were delivered in a ‘train’ structure with pulse trains at 10 Hz containing up to 352 pulses at an inter-train rate of 1.1 MHz.

The AGIPD-1M detector is an integrating detector capable of recording up to 352 diffraction images per train operating in a dynamic gain mode which lowers the gain for higher photon counts (Hantke *et al*., 2018). This allows both single photon detection and high dynamic range. The installed AGIPD-1M has 1024×1024 pixels of size 200×200 μm with a variable size central aperture For this set of trial experiments (proposal 2316) the detector was positioned 1.62 m from the sample—X-ray interaction point.

Diffraction patterns were collected during two days in 19 runs with 100 X-ray pulses per train. Total acquisition time was 2 hours 25 minutes and 8669500 images were recorded. We have identified 3800 images with scattering above background (hits) using the lit pixel counter (Hantke *et al*., 2014). Thus, hit rate was 0.044%. In order to filter out scattering from the salt balls, other non-repeatable particles or from multiple particles we have clustered radial profiles using hierarchical agglomerative clustering. Eliminating debris scattering clusters we finally got 750 diffraction patterns similar to the scattering from icosahedral virions with visible fringes. In figure 9A-D we show four representative hits form particles with slightly different sizes. Particle sizes were estimated by the fitting spherical form factor to the radial profiles of each diffraction pattern. Size distribution across all hits selected with cluster analysis is shown in the Figure 9E. Unfortunately, the size distribution is quite wide. It may indicate the aggregation of virion particles with buffer salt after evaporation of droplets.

**Figure 9.**
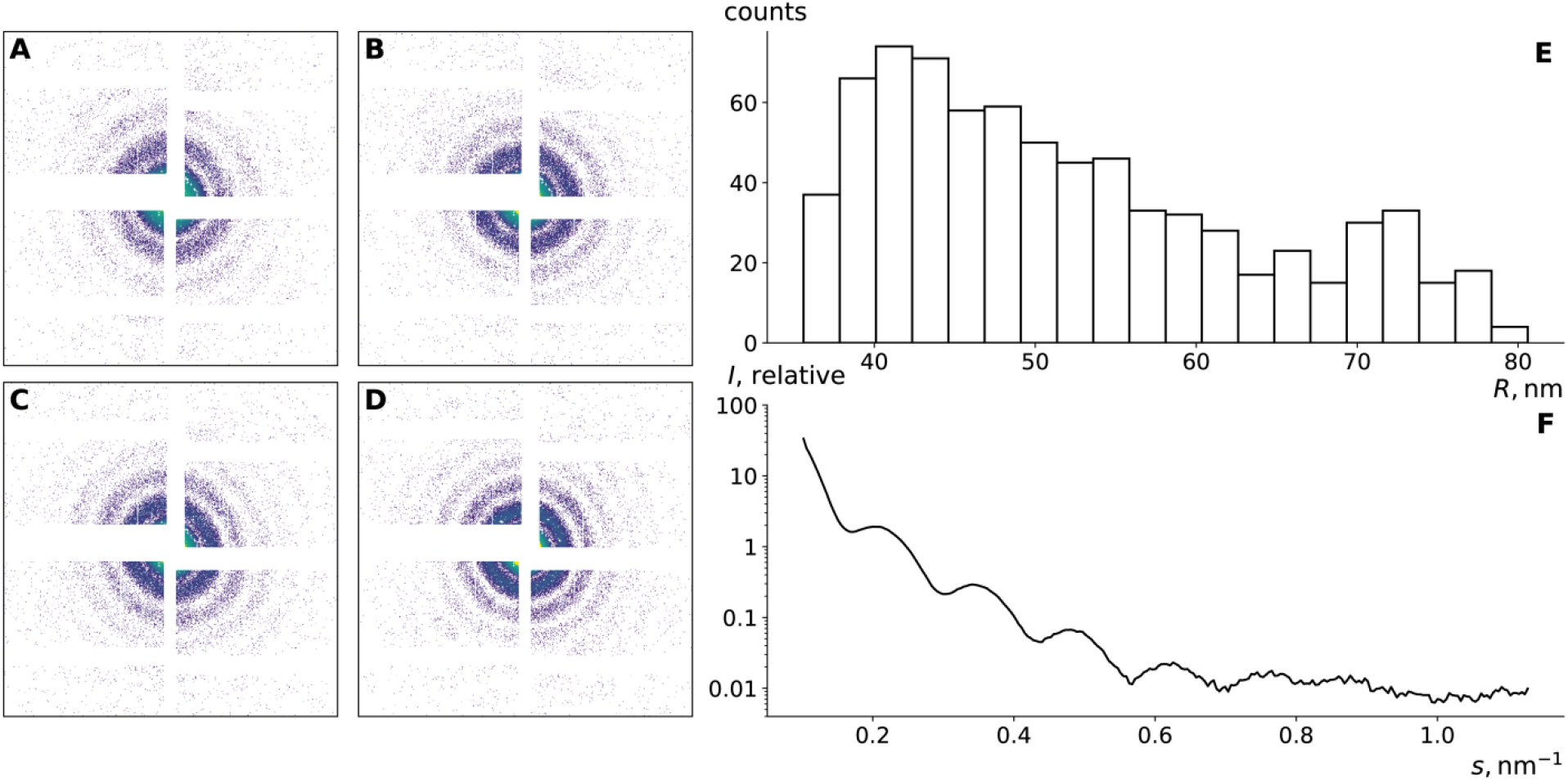
Diffraction patterns of iTBEV particles: (**A-D)** representative single particle diffraction patterns from particles with sizes 40.4, 45.9, 50.7, and 54.7 nm, correspondingly; (**E)** distribution of particle sizes estimated by fitting spherical form factor to the radial profiles of diffraction patterns; (**F)** radial profile averaged over similar diffraction patterns with mean size of 53.8 nm.

In Figure 9F we show the radial profile averaged in one of the clusters contained 125 images with scattering from samples with mean size of 53.8 nm and dispersion of 2.1 nm. On this profile at least five fringes are clearly visible despite the size distribution and fluctuations of the diffraction center, which allows us to conclude that the quality of the diffraction patterns is quite high.

### 3.6 iTBEV Sample Storage Influences Its Quality as Revealed by SAXS

The experimental scattering curves from inactivated TBEV particles prepared using the ultracentrifugation (protocol 3) immediately before the experiment (See section 3.2.2.) and after 5 months storage at −80 °C were compared (Table 1). Experimental SAXS curves from freshly prepared and frozen iTBEV particles and the corresponding distance distribution functions are shown in Fig. 5B. The radius of gyration and the maximal diameter of the particles in the recovered sample significantly increased relatively to the freshly prepared iTBEV particles, indicating the degradation of the sample. Evaluation of volume size distribution within the approximation of polydisperse spheres showed that the particles stored for 5 months were a mixture of spherical particles of 25 nm (90%) and 45 nm (10%) radius in solution (Table 1). This observation highlighted the importance of shipping the freshly prepared and unfrozen samples for SPI experiments, because during freeze-thaw cycles aggregation of the particles may occur in the system, leading to a substantial polydispersity of the particle size.

## 4. Discussion

Single-particle techniques offer unique opportunity to understand the role of structural variability in biological function (Ourmazd, 2019). For enveloped viruses, such as flaviviruses, cryo-EM and single particle imaging at XFEL are the only suitable ways to resolve the structure at the atomic level. Quality of the sample is a crucial milestone of any structural method. Requirements for the sample are similar for both single-particle imaging techniques, however, small sample volume is the feature of cryo-EM, whereas for SPI at XFELs a large sample volume is crucial to collect a sufficient amount of data. High concentration and homogeneity is necessary both for cryo-EM single particle analysis and SPI experiment at XFELs. Advanced computational classification approaches are developed for the cryo-EM data analysis, allowing to successfully “purify” sample *in silico* (Lyumkis, 2019) or even solve the structure of one protein from the mixture with another one (Pichkur *et al*., 2021). In SPI experiments, sample-sized impurities may produce parasitic scattering and along with aggregates may reduce target concentration in the sample considerably. Aggregation of the particles may also lead to capillary clogging during the injection. Cryo-EM is also more tolerant to buffer composition. Electrospray sample injection is usually performed in volatile buffers (mainly ammonium acetate) with conductivity in the range of 1.5-4 mSm/cm, which is considered sufficient for sample mobility. Thus, more strict requirements to the sample amount and quality for SPI experiments at XFELs highlight the need for development of advanced protocols for virus sample preparation.

In this work an enveloped virus was tested for the first time as an object for SPI at free electron laser, and principal possibility to collect single particle diffraction data was demonstrated. Physiological buffer was used for injection. For this structural method, several mL of highly purified homogenous virus suspension with concentration 10^12^-10^13^ particles/mL is needed. Such a demand may be fulfilled at the factories for inactivated vaccine production. Particularly, strain Sofjin of the Far-Eastern subtype is used for manufacturing of a commercial inactivated vaccine against TBEV. This strain was chosen as a model for the current study. Formaldehyde-inactivated TBEV virions retain E protein antigenic sites that was confirmed by ELISA with the E protein mAbs and Western-blot with rabbit polyclonal antibodies or mAbs. In addition, the method of TBE virus inactivation by formaldehyde are successfully used for more than 60 years for obtaining highly immunogenic and protective TBE vaccines (Ruzek et al., 2019). Combination of purification techniques accompanied by standard biochemical methods for estimation of infectivity, concentration, etc., and quality control pipeline mainly based on TEM and SAXS, allowed us to prepare a concentrated, homogenous, non-aggregated TBEV sample. Complementary physical methods of analysis are preferable for different steps of the sample production pipeline. Both TEM approaches helped to indicate sample heterogeneity during the ion exchange chromatography step. Cryo-EM helped to verify samples’ homogeneity. SAXS was useful at the final step of protocol choice. After the sample quality control by TEM and SAXS, protocol 3 gave the most consistent, homogeneous, pure, and concentrated suspension of viral particles, and was then used for a large-scale sample preparation for an SPI experiments at the European XFEL. Although storage of viruses at cryogenic temperatures is a common practice in structural virology, our SAXS investigation pinpointed importance of using fresh sample for injection and data collection during SPI at XFELs.

We demonstrated a diffraction pattern obtained with a single X-ray pulse from iTBEV particle for the first time — using SPB/SFX instrument of the European XFEL. Achievement of diffraction from such a small single viral particle, with diameter of approx. 50 nm, shows possibilities of instrument. Our purification and quality assessment pipeline may be useful for the preparation of other challenging virus samples for single particle imaging at different FELs.

## Acknowledgements

We thank Ekaterina Round, Robin Schubert for help with sample characterization at XBI laboratory, XFEL as well as Johan Bielecki, Richard Bean, Egor Sobolev, Adrian P. Mancuso, Mikhail Rychev and Serguei Molodtsov for help with SPI experiment and data processing. We acknowledge the European XFEL in Schenefeld, Germany for the provision of X-ray free-electron laser beamtime on the SPB/SFX scientific instrument and would like to thank the staff for their assistance.

