## Supporting Information for "Preparation and Characterization of Inactivated Tick-Borne Encephalitis Virus Samples for Single Particle Imaging at European XFEL"

$ These authors contributed equally

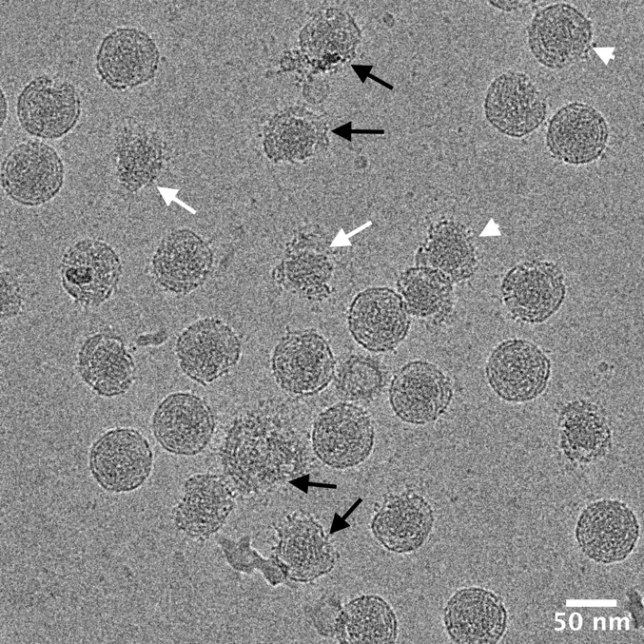

**Figure S1.** Cryo-electron microscopy of iTBEV. The sample contained mature particles, immature particles (white arrows), broken particles (black arrows), and semi-mature particles (white arrowheads).

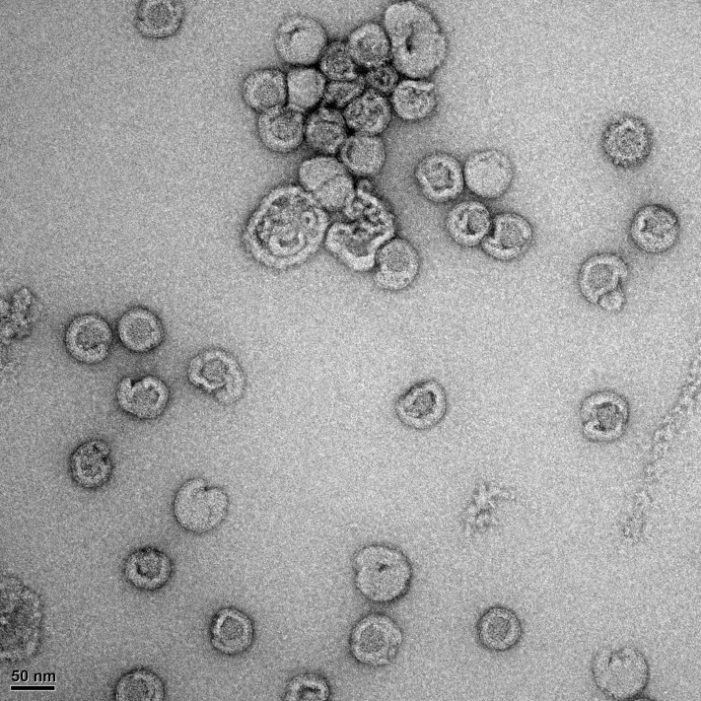

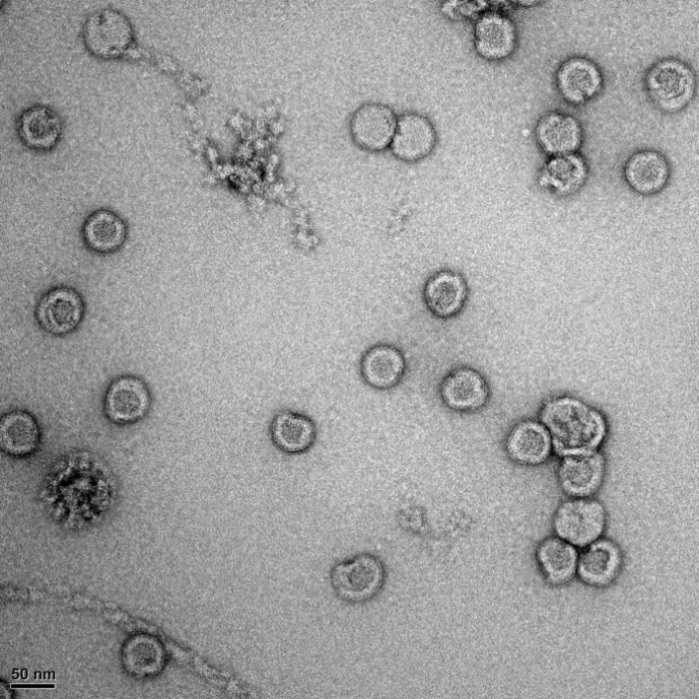

**Figure S2.** Negative stain TEM of iTBEV samples prepared using Protocol 1(Figure 2, main text)

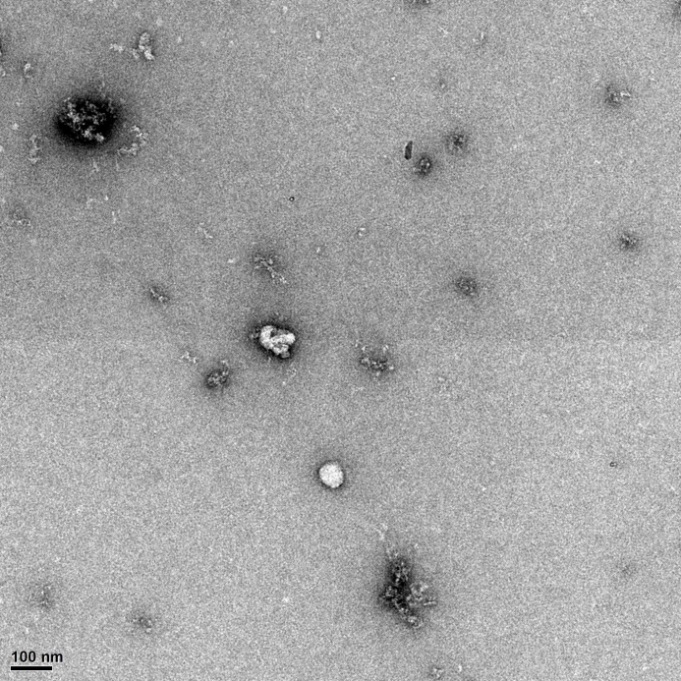

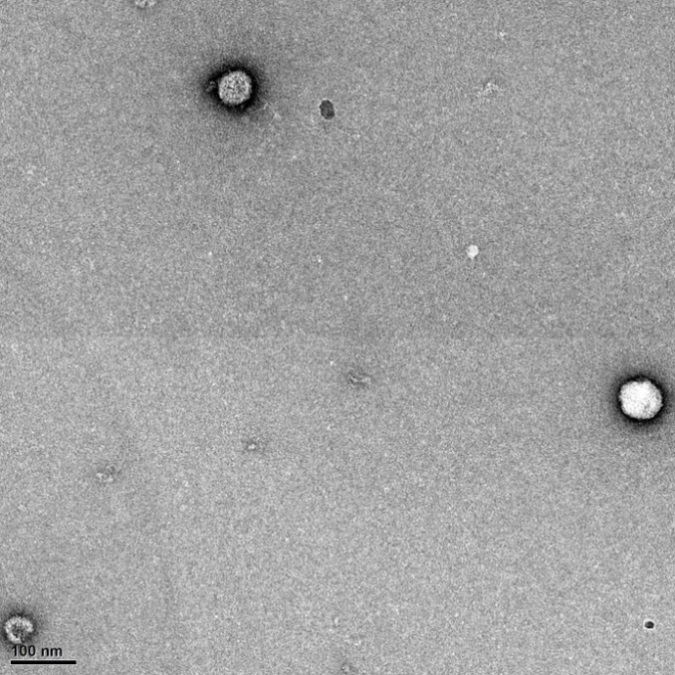

**Figure S3.** Negative stain TEM of iTBEV prepared using Protocol 2. (Figure 2, main text)

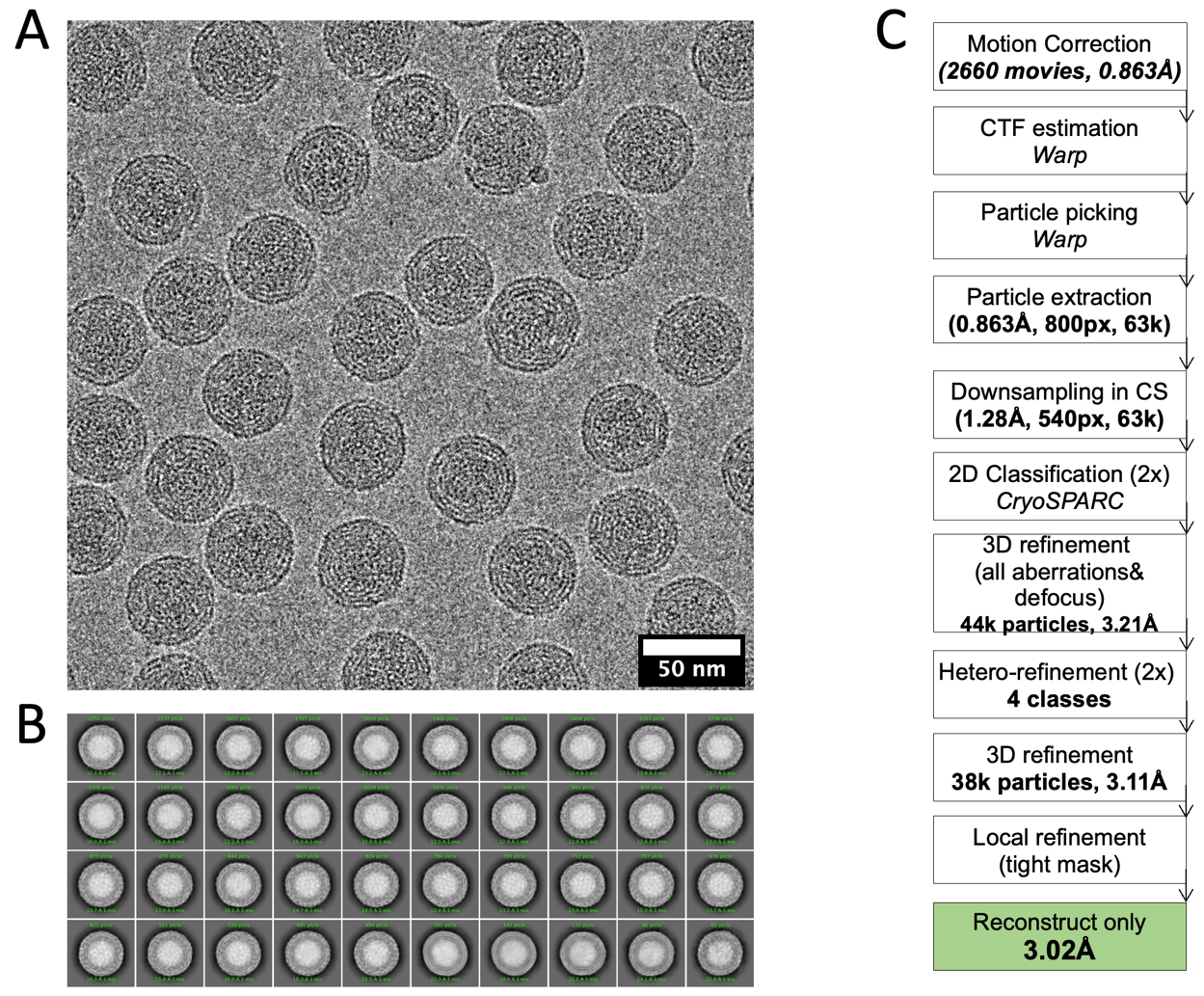

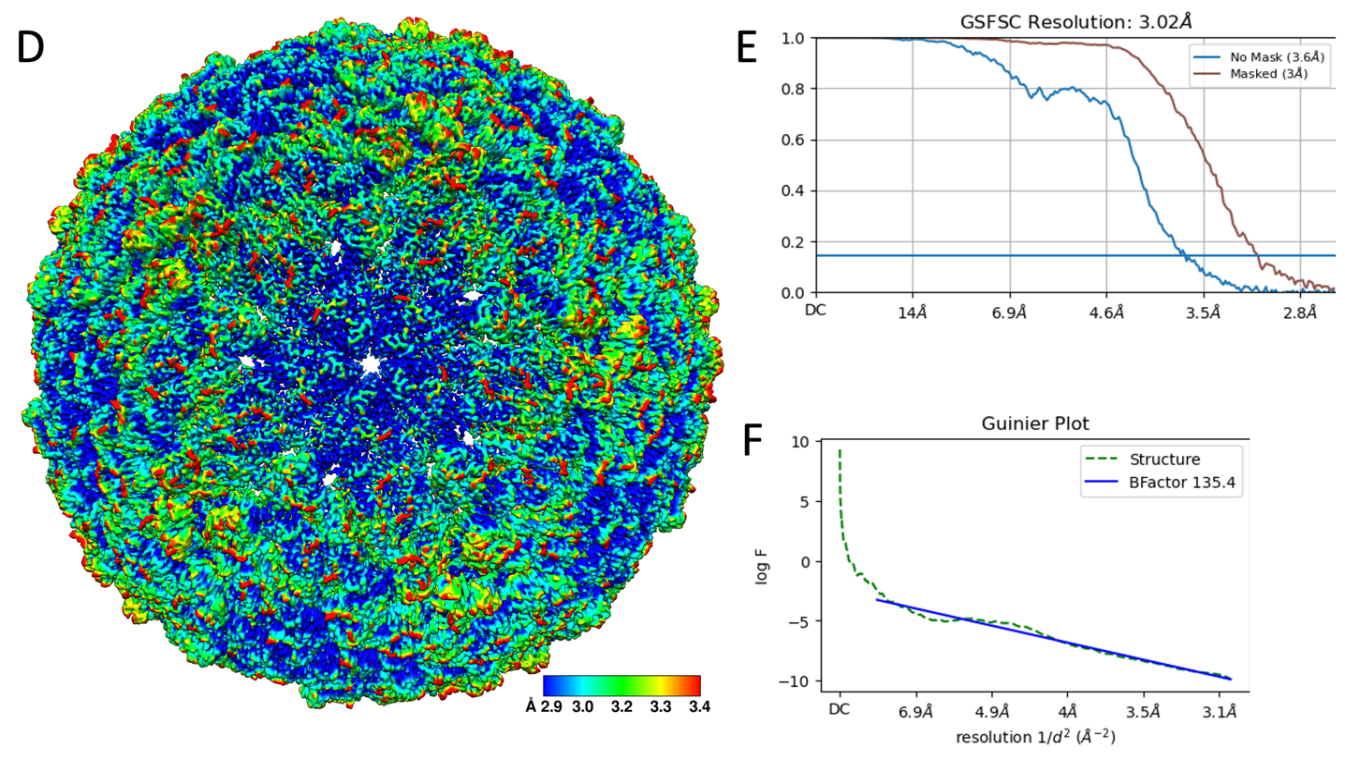

**Figure S4.** Cryo-EM data processing workflow. (A) Representative experimental cryo-EM image (B) Representative 2D classes after second run of 3D classification (C) Data-processing workflow. (D) Quality assessment of the cryo-EM data, local resolution map, calculated in cryoSPARC with the FSC=0.5 criteria (E) FSC plot (F) Guinier plot.
